# A role for MCH neuron firing in hippocampal plasticity and learning

**DOI:** 10.1101/2022.12.01.518339

**Authors:** Julia J. Harris, Cristina Concetti, Daria Peleg-Raibstein, Denis Burdakov

## Abstract

It has been revealed that melanin-concentrating hormone (MCH) neurons in the hypothalamus can influence learning (Liu et al., 2022) and memory formation (Kosse & Burdakov, 2019), but the cellular mechanisms by which they perform this function are not understood. Here, we examine the role of MCH neural input to the hippocampus, and show *in vitro* that optogenetically increasing MCH axon activity facilitates hippocampal plasticity by lowering the threshold for synaptic potentiation. *In vivo*, we find that MCH neurons are naturally active in response to reinforcing cues during a spatial learning task, and that this activity is correlated with the speed of learning. Together, our results align with increasing evidence that MCH neurons play an ‘on-line’ regulatory role in learning, and reveal that this could be achieved through modulation of synaptic plasticity in the hippocampus.

## INTRODUCTION

Synaptic plasticity in the hippocampus is a major cellular substrate for learning, but the plastic changes that occur there do not happen in isolation. The hippocampus receives a variety of input projections from different brain regions, which could theoretically modulate synaptic plasticity during learning, according to their unique environmental responses. One such candidate are hypothalamic neurons containing the neuropeptide transmitter, melanin-concentrating hormone (MCH). MCH neurons project widely from the lateral hypothalamus (LH) throughout the brain (Bittencourt et al., 1992, 2011; Steininger et al., 2004; Arrigoni et al., 2019), with particularly dense innervation of the hippocampus (Izawa et al., 2019). Infusion of the MCH peptide itself directly into the hippocampus improves memory retention (Monzon et al., 2009), and silencing MCH neurons during an object recognition task disrupts memory formation (Kosse & Burdakov, 2019). *In vitro* work has shown that when MCH neurons (Le Barillier et al., 2015) or MCH receptors (Pachoud et al., 2010) are genetically deleted, hippocampal plasticity is impaired such that a larger stimulus is required to induce long term potentiation (LTP).

These lines of evidence suggest that MCH input to the hippocampus may be capable of modulating synaptic plasticity, and would predict that *increased* MCH input should *facilitate* synaptic plasticity. Here, we tested this hypothesis directly, by optogenetically activating MCH axons in hippocampal slices. We found that increased MCH activity in the hippocampus lowers the threshold for lasting synaptic potentiation. We then examined whether MCH neurons would be able to perform such a role *in vivo*, by studying their natural activity during a hippocampal learning task. Using fibre photometry, we found that MCH neurons increased their activity in response to reinforcing cues during a spatial learning task, and that this activity correlated with the speed of learning.

Our results provide evidence that MCH neuron activity in the hippocampus can facilitate local synaptic plasticity, and also show that such neural activity is naturally modulated during a spatial learning task. Together, these findings support the idea that MCH neurons may regulate spatial learning, through modulatory effects on hippocampal synaptic plasticity.

## RESULTS

We stereotaxically injected cre-inducible channelrhodopsin (ChR2(H134R)-EYFP) bilaterally into the mouse lateral hypothalamus of MCH-Cre mice (Figure 1A,B). Whole-cell patch-clamping confirmed that all EYFP-expressing cells tested (11/11) responded to blue light stimulation (Figure 1C).

**Figure 1.**
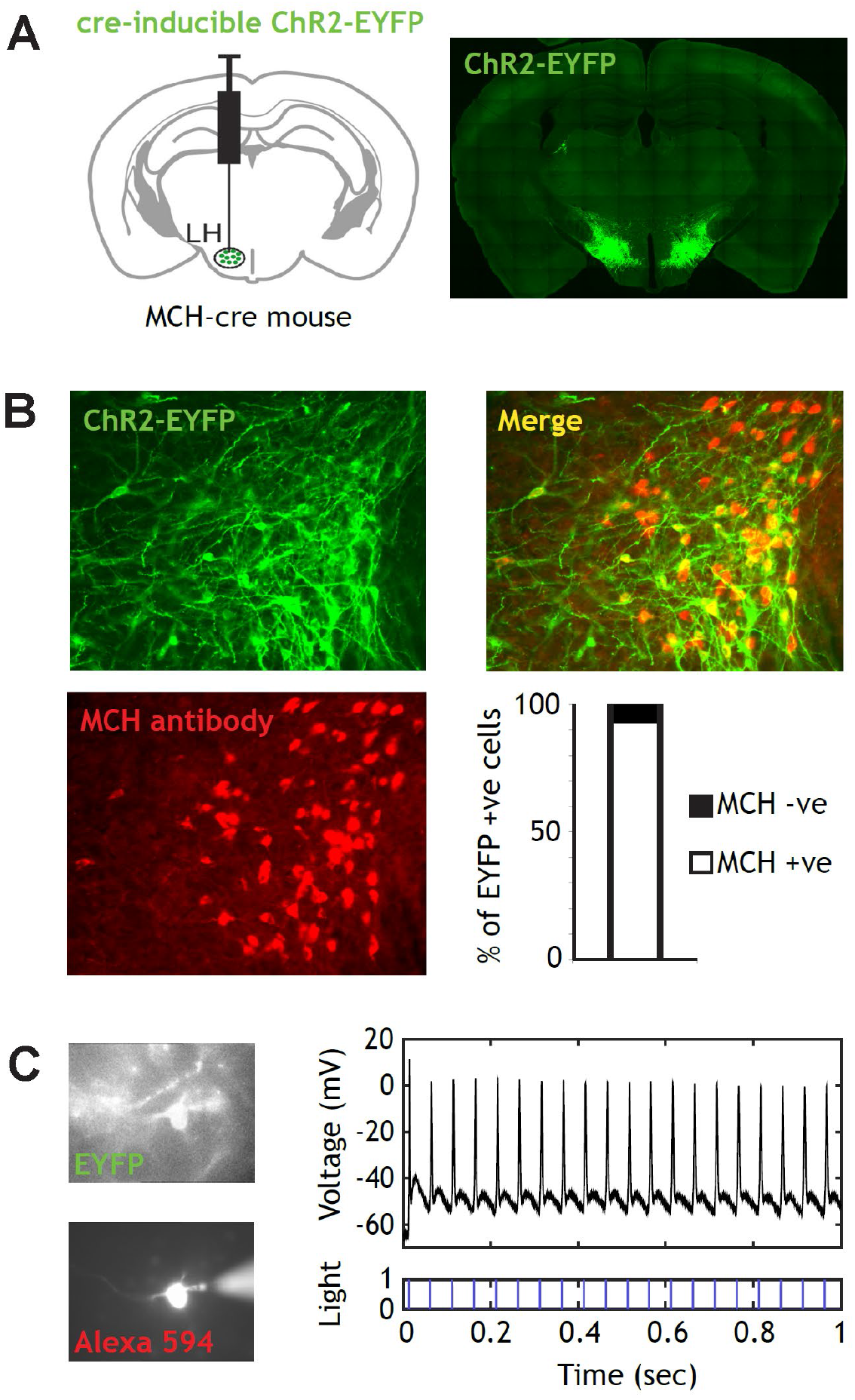
Channelrhodopsin expression in hypothalamic MCH neurons. (**A**) Bilatereral stereotaxic injection of cre-inducible ChR2(H134R)-EYFP into the lateral hypothalamus (LH) of MCH-cre mice. (**B**) ChR2-EYFP is expressed in cell bodies and axons of neurons in the LH. 93% (311/336) of cell bodies expressing ChR2-EYFP also stain for MCH antibody. (**C**) Whole cell patch clamping in the LH. 100% (11/11) of EYFP-expressing cells fire action potentials in response to blue light. 0% (0/9) of non-fluorescent cells respond to blue light (not shown).

Whole-cell patch-clamping of hippocampal pyramidal neurons did not reveal direct synaptic input from MCH axons (0/18 pyramidal neurons showed time-locked responses to blue illumination of MCH-ChR2 axons in the hippocampus, Figure 2A). Optogenetically activating MCH axons (30 sec stimulation at 20 Hz, designed to trigger peptide release, Schöne et al., 2014) did not affect the frequency of spontaneous excitatory postsynaptic currents (sEPSCs) or spontaneous inhibitory postsynaptic currents (sIPSCs) recorded in pyramidal cells, nor the amplitude of sIPSCs (Figure 2B). However, sEPSC amplitude was significantly reduced following blue light, in MCH-ChR2+ compared to MCH-ChR2-brain slices (Figure 2B).

**Figure 2.**
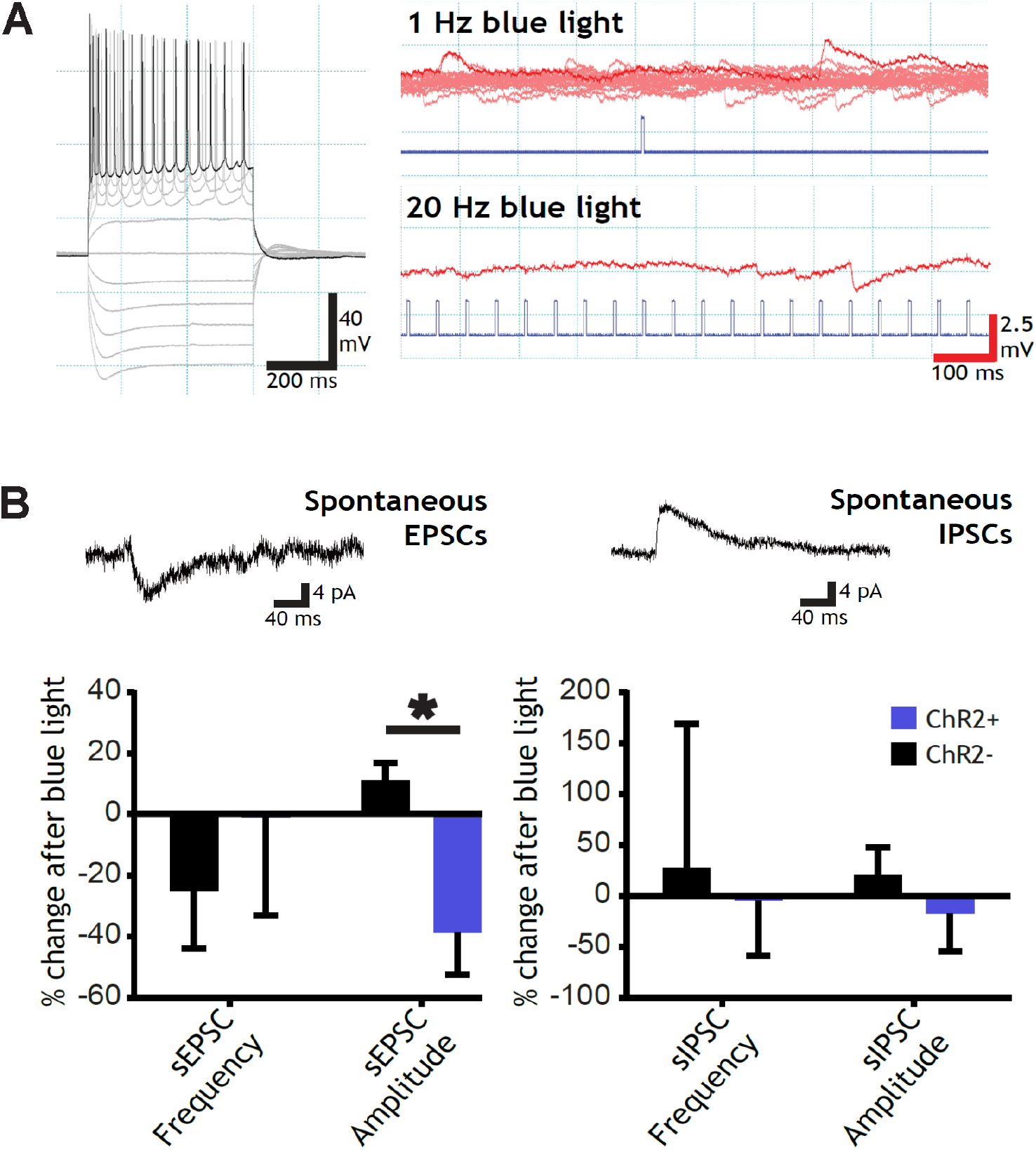
MCH axon activation reduces spontaneous excitatory synaptic current (sEPSC) size in hippocampal pyramidal neurons. (A) Whole cell patch clamping in the hippocampus. Hippocampal pyramidal neurons do not receive direct synaptic input from MCH axons (0/18 pyramidal cells show time-locked responses to blue light in MCH-ChR2 slices, example right). (B) Optogenetically activating MCH axons (20 Hz for 30 sec) did not affect the frequency of sEPSCs (H-B adjusted p = 0.52) or sIPSCs (H-B adjusted p = 0.61) recorded in pyramidal cells of MCH-ChR2+ slices compared to MCH-ChR2-slices. sIPSC amplitude was also unaffected (H-B adjusted p = 0.19), but sEPSC amplitude was significantly reduced in the three minutes following blue light, in MCH-ChR2+ compared to MCH-ChR2-brain slices (Holm-Bonferroni adjusted p=0.017).

We next examined the effects of optogenetic activation of MCH axons in the hippocampus on the plasticity of pyramidal cell excitatory field potentials (fEPSPs, Figure 3A,B). We implemented two classical potentiation protocols: a weak potentiating stimulus (1 tetanus; typically inducing brief post-tetanic potentiation) and a strong potentiating stimulus (four tetani; typically inducing LTP), each interleaved with blue light stimulation of MCH axons (Figure 3C,D left).

**Figure 3.**
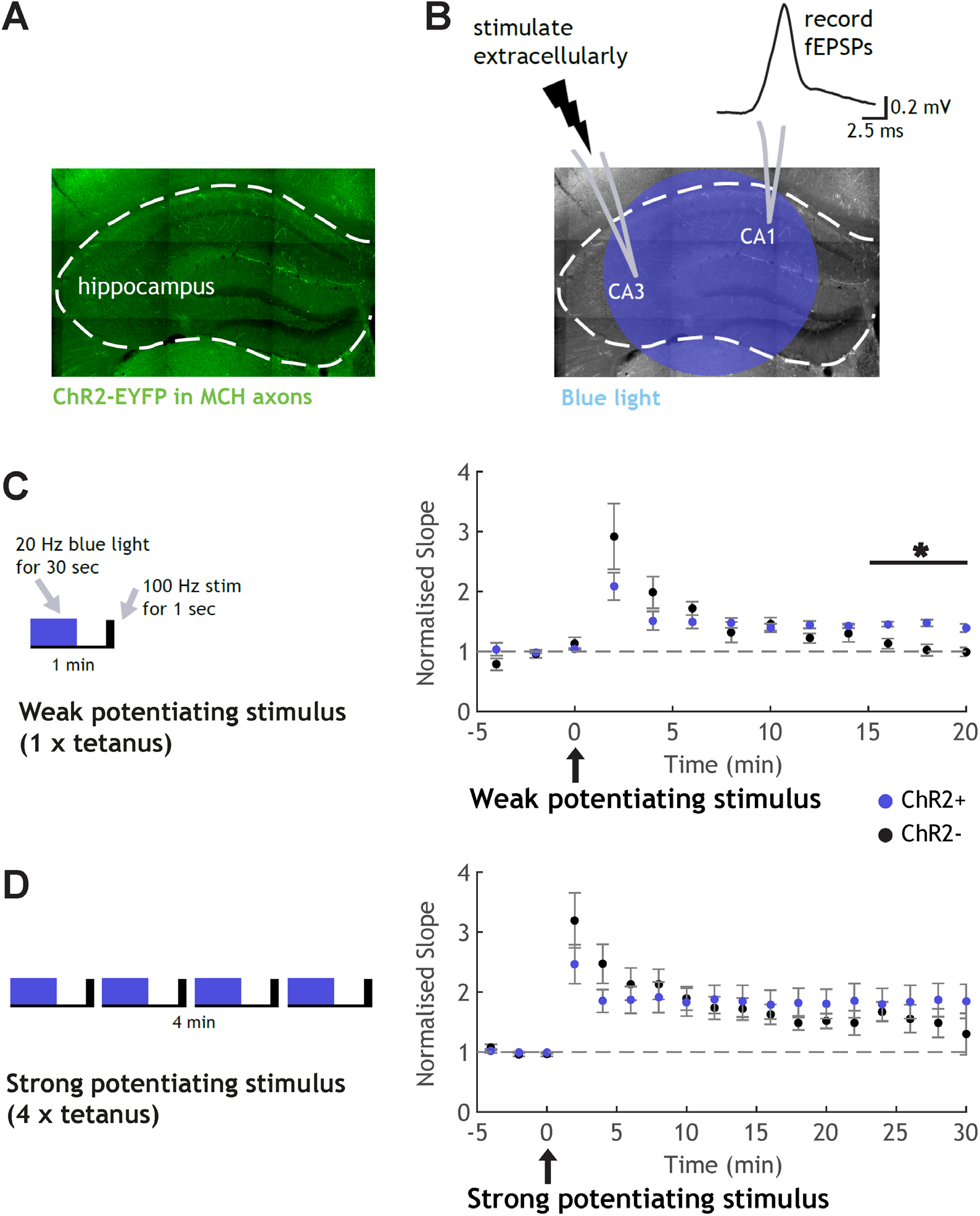
MCH axon activation lowers threshold for synaptic potentiation. (A) ChR2-EYFP-expressing MCH axons were visible in the hippocampus. (B) Excitatory postsynaptic field potentials (fEPSPs) were generated by extracellular stimulation in CA3 and recorded in CA1 (top right). (C,D left) Electrical stimulation patterns designed to trigger two different types of classical synaptic potentiation were combined with blue light illumination of the hippocampus to active MCH axons. (C, right) Optogenetic activation of MCH axons in the hippocampus altered the synaptic plasticity response to a weak potentiating stimulus (two-way ANOVA interaction effect: F(9, 110) = 2.592, p = 0.0095). In MCH-ChR2-control slices, a weak potentiating stimulus led to “post-tetanic potentiation”, which is short-term, with fEPSPs decaying back to baseline levels by ~15 mins (black points). In MCH-ChR2+ slices, the same electrical and optogenetic stimulation led to longer lasting potentiation (blue points), which was significantly greater than in control slices at 15-20 minutes post-induction (Holm-Bonferroni adjusted p = 0.005). (D, right) Optogenetic activation of MCH axons in the hippocampus did not significantly alter the response to a strong potentiating stimulus (two-way ANOVA, no significant interaction: F(14, 165) = 0.9, p = 0.5584).

Optogenetic activation of MCH axons in the hippocampus significantly altered the plasticity response to the weak potentiating stimulus (Figure 3C; two-way ANOVA: F(9, 110) = 2.592, p = 0.0095): In MCH-ChR2-control slices, one electrical tetanus plus light led to short-term potentiation, with fEPSPs decaying back to baseline by ~15 mins (black points). In MCH-ChR2+ slices (blue points), the same electrical and optogenetic stimulation led to potentiation that did not decay after 10 minutes, and remained significantly elevated compared to control slices 15-20 minutes post-induction (Holm-Bonferroni adjusted p = 0.005).

On the other hand, optogenetic activation of MCH axons in the hippocampus did not significantly alter the response to the strong potentiating stimulus (Figure 3D; two-way ANOVA revealed no significant interaction, p > 0.05), nor to a strong depressing stimulus (900 single electrical pulses delivered at 1 Hz; Supplementary Figure 1; two-way ANOVA revealed no significant interaction, p > 0.05).

These results suggest that MCH activity can facilitate hippocampal plasticity by lowering the threshold for synaptic potentiation.

We next examined whether MCH neurons would be able to perform such a role *in vivo*, by studying their natural activity during a hippocampal learning task. Mice expressing GCaMP6s in the lateral hypothalamus under the control of the MCH promoter were implanted with an optic fiber for *in vivo* imaging of calcium fluorescence (Figure 4A). They then performed trials in a T-maze, learning which arm contained a food reward and which contained an empty food cup (Figure 4B). While we noticed an increase in GCaMP6s fluorescence when mice entered the food cup zone in both correct (two-tailed, one-sample t-test, p value < 0.001) and incorrect (two-tailed, one-sample t-test, p value < 0.001) trials, this increase was significantly larger for correct trials (two-tailed, unpaired t-test, p value = 0.0021, Bonferroni-adjusted alpha = 0.0167; Figure 4C). Interestingly, mice with larger GCaMP6s responses during correct trials tended to learn the task faster than mice with smaller GCaMP6s responses, requiring fewer trials to reach criterion performance (Figure 4D, left). No such trend was found for incorrect trials (Figure 4D, right). These results suggest that MCH activity in response to positively reinforcing cues could act as a learning signal during hippocampus-dependent memory acquisition.

**Figure 4.**
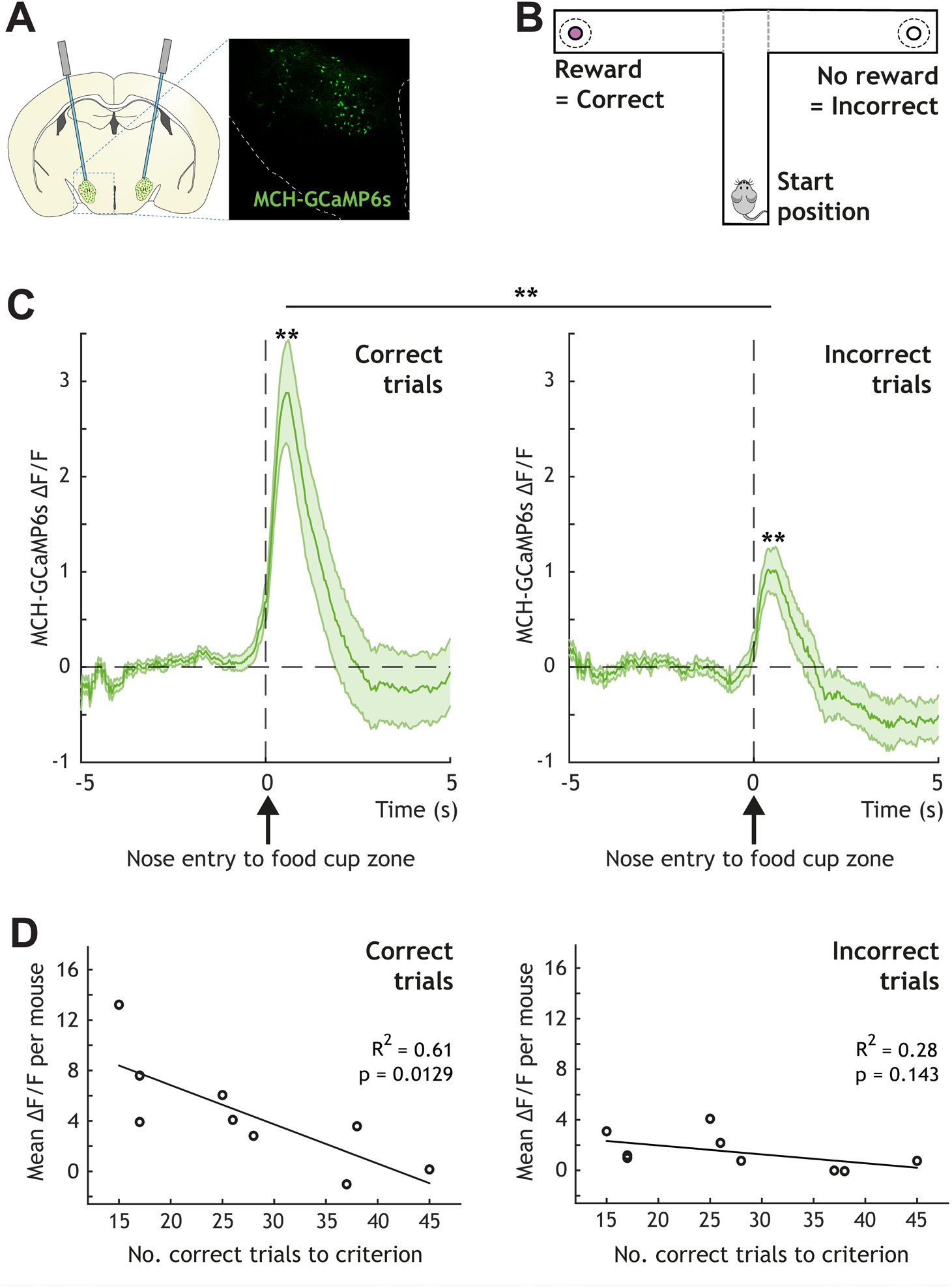
MCH neuron activity increases during reward discovery and correlates with speed of learning. (A) GCaMP6s was expressed in the lateral hypothalamus (LH) under the control of the MCH promoter, and fluorescence was monitored *in vivo* using fiber photometry. (B) GCaMP6s fluorescence was recorded while mice performed trials in a T-maze. At the end of each arm was a food cup (solid circle) but only one arm contained a liquid food reward (strawberry milkshake - purple centre). The position of the food reward was kept the same for each mouse. A 3 cm perimeter around each food cup was defined as the “food cup zone” (dashed circle). (C) MCH-GCaMP6s fluorescence was significantly elevated at food-cup-zone entry (time = 0), for both correct (two-tailed, one-sample t-test, p value < 0.001) and incorrect (two-tailed, one-sample t-test, p value < 0.001) trials, but significantly more for correct trials (two-tailed, unpaired t-test, p value = 0.0021) (averaged trials from 9 mice - adjusted alpha = 0.0167 with Bonferroni correction). (D) Across mice, the amount of MCH-GCaMP6s fluorescence during correct trials was negatively correlated with the number of correct trials needed to achieve criterion performance. This trend was not present for MCH-GCaMP6s fluorescence during incorrect trials. Each data point is the averaged amplitude (across trials, across session) of MCH-GCaMP6s fluorescence upon entry of the nose point into the food-cup zone for one mouse (n = 9 mice).

## DISCUSSION

We have found that optogenetically activating MCH axons in the hippocampus facilitates synaptic plasticity *in vitro*, by lowering the threshold for lasting potentiation induced by electrical stimulation. *In vivo*, we discovered that MCH neurons become naturally active in response to reinforcing cues, and that this activity is correlated with learning in a hippocampus-dependent task. These two lines of evidence suggest that MCH neurons could provide a valuable learning signal, lowering the threshold for hippocampal plasticity in order to facilitate learning in response to salient cues.

Previous studies have shown that abolishing MCH activity in the brain impairs hippocampal plasticity and learning: in mice congenitally lacking MCH receptors, the threshold for hippocampal plasticity is increased (Pachoud et al., 2010), and in mice with selective deletion of MCH neurons in adulthood, hippocampal post-tetanic potentiation is diminished and short-term memory is disrupted (Le Barillier et al., 2015). These works suggest that MCH input to the hippocampus may be necessary for normal hippocampal plasticity, but as with all genetic knockout approaches, it is hard to rule out the possibility that compensatory mechanisms are responsible for the effects. Here, we show in intact circuits that *increasing* MCH axon activation in the hippocampus *reduces* the threshold for plasticity induction and *increases* the longevity of post-tetanic potentiation. This evidence supports the idea that MCH input to the hippocampus can modulate hippocampal plasticity. Whether this is likely to be operating *in vivo* depends on whether MCH neurons are actually active during learning.

Previous work from the lab showed that MCH neurons are indeed active during object learning, and that this activity is crucial for the formation of new object memories (Kosse & Burdakov, 2019). Here, we examined whether these neurons are also active during a spatial learning task. Using *in vivo* calcium imaging in naturally behaving mice, we found that MCH neuron activity increased in response to reinforcing cues, and that the extent of this natural activity was correlated with the speed of learning. Infusion of the MCH peptide directly into the hippocampus has also been shown to increase learning in a step-down avoidance task (Monzon et al., 1999).

These results all point towards the possibility of MCH neurons performing a regulatory role during learning, underpinned by modulatory effects on local synaptic plasticity. In line with this hypothesis, MCH application in the hippocampus has been shown to directly augment synaptic transmission (Varas et al., 2002) and, very recently, MCH projections to the dorsolateral septum have been revealed to increase the efficacy of its hippocampal inputs, ultimately facilitating spatial learning (Liu et al., 2022).

Overall it is becoming clear that MCH neurons in the lateral hypothalamus play a vital role in modulating learning and memory formation, through their direct responses to the environment and their projections to many brain regions, where they have the capacity to directly alter synaptic transmission and plasticity. The importance of MCH neurons in cognition is underlined by the memory-preserving effect of MCH peptide found in mouse models of Alzheimer’s disease (Oh et al., 2019).

One interesting feature of MCH neurons is that they are highly active during REM sleep (Hassani et al., 2009). Given the many lines of evidence discussed above which suggest that MCH neurons augment synaptic potentiation and learning, and an entirely different body of literature which suggests that REM sleep is important for memory (reviewed in Wamsley and Stickgold, 2011), it would be sensible to guess that MCH neurons contribute to this night-time consolidation. It was therefore surprising when Izawa et al. (2019) recently revealed apparently the opposite: that MCH activity during REM sleep aids *forgetting*. Importantly, Izawa and colleagues also found evidence that wake-active and REM-active MCH neurons are distinct subsets within the hypothalamus. These subsets could modulate hippocampal plasticity in different ways. For instance, the REM-active MCH neurons were found to increase IPSC occurrence in the hippocampus, while the MCH neurons that we examined reduced EPSC amplitude, suggesting effects on GABAergic and glutamatergic inputs, respectively. We would predict wake-active MCH neurons to facilitate potentiation, perhaps through effects on glutamatergic transmission, and REM-active MCH neurons to either inhibit potentiation or facilitate depression, perhaps through effects on GABAergic transmission. Alternatively, it is possible that MCH neurons do not drive remembering or forgetting per se, but instead play a permissive role by acting as “eligibility traces” for plasticity, whereupon other inputs can potentiate or depotentiate synaptic connections (Burdakov & Peleg-Raibstein, 2020).

To understand the full picture of how MCH neurons contribute to remembering and forgetting, it will be essential to examine how their activity modulates the cellular mechanisms of plasticity, across different vigilance states and in a sub-population-specific manner. By lowering the threshold for lasting potentiation, the results presented in this paper contribute a potential mechanism by which some of these neurons could aid memory formation during learning.

---

**Supplementary Figure 1.**
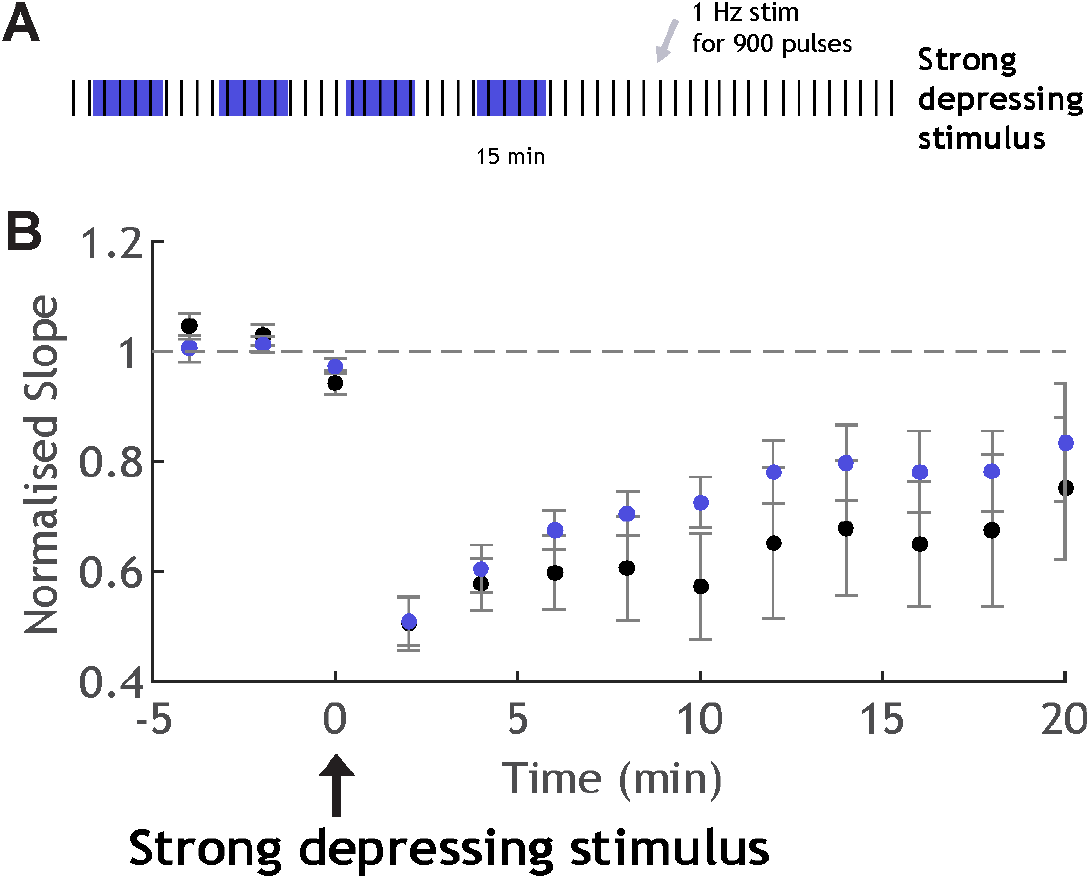
No effect on synaptic depression. (A) Electrical stimulation pattern designed to trigger synaptic depression (900 electrical pulses delivered once per second) was combined with blue light illumination of the hippocampus to activate MCH axons (20 Hz for 30 seconds delivered four times). (B) This optogenetic activation of MCH axons in the hippocampus did not significantly alter the response to the strong depressing electrical stimulation (two-way ANOVA, no significant interaction: F(9, 110) = 1.21, p = 0.2945).

**Supplementary Figure 2.**
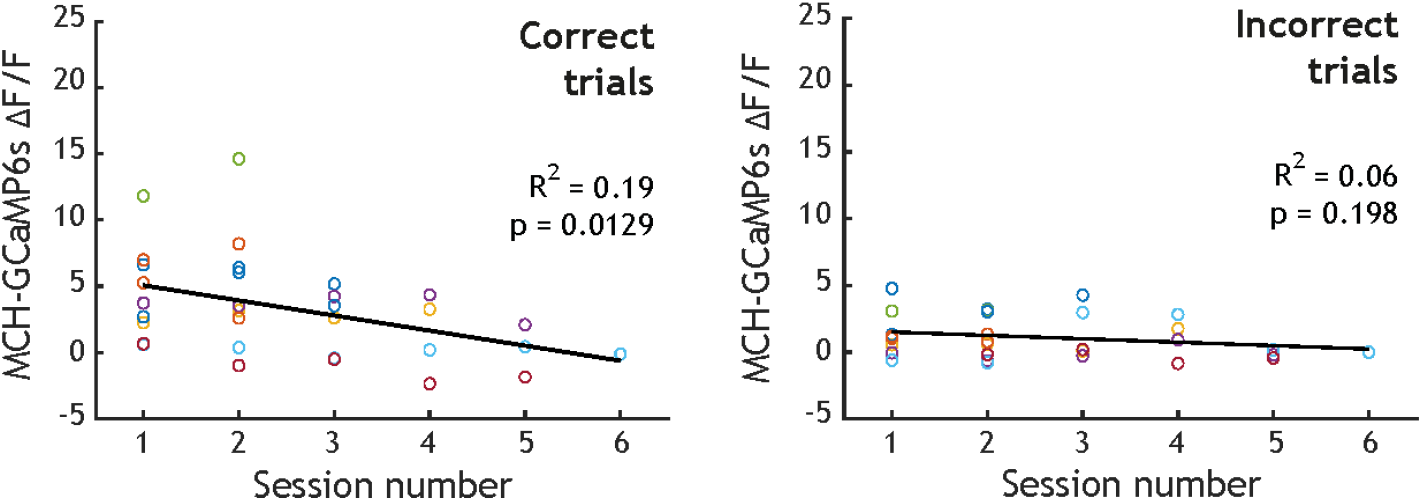
MCH activity for individual mice across sessions. For correct trials, the food cup entry-associated MCH activity tended to decrease as sessions progressed, whereas no such trend was present for incorrect trials.

## METHODS

### Animals

Animal research has been approved by the United Kingdom Home Office and the Animal Welfare and Ethical Review Panel of the Francis Crick Institute, and Zurich Cantonal Veterinary Office. All procedures were performed in accordance with the Animals (Scientific Procedure) Act of 1986 (UK) and the Animal Welfare Ordinance (TSchV 455.1) of the Swiss Federal Food Safety and Veterinary Office (Switzerland). Mice were kept on a standard chow and water ad libitum and on a 12-h/12-h light/dark cycle. Behaviour experiments were performed during the dark phase, using males and females at least 8 weeks of age. Injections for slice physiology experiments were performed on males and females at least 4 weeks of age, and slices were obtained 14-28 days later.

### Genetic Targeting

The specific targeting of GCaMP6s to MCH neurons was performed using the same genetic tools as described and histologically validated in our previous studies (Kosse & Burdakov, 2019; Concetti et al., 2020). In brief, we injected an AAV vector carrying the 0.9-kb preproMCH gene promoter AAV9.pMCH.GCaMP6s.hGH (1.78 × 1014 gc/mL; Vigene Bioscience, characterised to target MCH cells with >90% specificity in Kosse & Burdakov, 2019) into the lateral hypothalamus of C57BL6 mice.

To target the light-gated ion channel, ChR2 to MCH neurons, we injected Cre-dependent AAV1.EF1a.DIO.hChR2(H134R)-eYFP.WPRE.hGH (>1.4 × 1013 gc/ml; Penn Vector Core, as in Apergis-Schoute et al., 2015) bilaterally into the lateral hypothalamus of the previously characterised and validated MCH-Cre mice (Kong et al., 2010). Confirmation of functional ChR2 expression was performed using whole-cell patch clamping combined with photostimulation in acute brain slices (Figure 1).

For stereotaxic brain injections, mice were anaesthetised with isoflurane and injected with meloxicam (Metacam, 2 mg/kg of body weight, s.c.) for analgesia. In a stereotaxic frame (Kopf Instruments), a craniotomy was performed, and a 33-gauge needle mounted on a Hamilton syringe was used to inject AAV vectors into the hypothalamus. Three injections (each 50 nL, at a rate of 50 nL/min) were administered per hemisphere at the following coordinates: bregma, −1.35 mm; midline, ±0.90 mm; depth, 5.70 mm, 5.40 mm, and 5.10 mm (Gonzalez et al., 2016; Kosse & Burdakov, 2019; Concetti et al., 2020). Before the behaviour experiments, the mice were allowed to recover from surgery for at least 10 days. Before the slice experiments, ChR2 expression was allowed to develop for 14-28 days.

### Immunohistochemistry

50 um cryosections were were stained for MCH using the primary antibody, rabbit polyclonal MCH (1:2000; Phoenix Pharmaceuticals, H-070-47) and the secondary antibody, Alexa Fluor 555 anti-rabbit IgG (1:500; Invitrogen, A-21244). Slices were DAPI-stained and mounted on slides, and images were captured using a Nikon NIS microscope or a Zeiss Axioscan slide scanner.

### Slice Physiology

After cervical dislocation, the brain was rapidly removed and immersed in ice-cold, slicing solution containing (in mM) 87 NaCl, 25 NaHCO3, 7 MgCl2, 2.5 KCl, 1.25 NaH2PO4, 0.5 CaCl2, 25 glucose and 75 sucrose, saturated with 95% O2/5% CO2 (modified from Bischofberger, 2006; Harris et al, 2015). The brain was sectioned into 350 μm coronal slices while submerged in ice-cold continuously oxygenated slicing solution. Slices were placed in a storage chamber containing continuously oxygenated slicing solution at room temperature naturally. During the experiment, slices were continuously perfused with artificial cerebrospinal fluid (aCSF) containing (in mM) 124 NaCl, 26 NaHCO3, 1 glucose, 2.5 KCl, 2.5 CaCl2, 1 NaH2PO4, and 1 MgCl2. The aCSF was heated to 35°C and constantly bubbled with 95% O2/5% CO2.

MCH neurons in the lateral hypothalamus were selected for whole-cell recording by their expression of ChR2-EYFP, visualised using a customised filter set (excitation 510/10 nm, dichroic 520 nm, emission 542/27 nm, Laser 2000; Figure 1). Pyramidal cells in the hippocampus were selected according to the shape and size of the soma, using differential interference contrast optics (Olympus; Figure 2). Whole-cell recordings from MCH neurons in the LH and pyramidal neurons in the hippocampus were obtained using 2- to 3-MΩ borosilicate glass electrodes filled with internal solution containing (in mM) 130 K-gluconate, 10 EGTA, 10 HEPES, 4 NaCl, 4 MgATP, 1 CaCl2, 0.5 Na2GTP, and Alexa Fluor 594 dye.

Recordings were made with a HEKA EPC10 USB Patch Clamp Amplifier, filtered at 5 kHz, and sampled at 20 kHz, and data was acquired using the Patchmaster software system (HEKA Electronik). Spontaneous intracellular currents (Figure 2) were analysed using Minianalysis (Synaptosoft Software).

### Plasticity Protocols and Photostimulation

For plasticity experiments, slices containing intact CA3 and CA1 regions of the hippocampus were selected, and the presence of EYFP-expressing MCH fibers in the hippocampus was confirmed (Figure 3A). Excitatory postsynaptic field potentials (fEPSPs) were evoked in the stratum radiatum of CA3 using current pulses delivered via a concentric bipolar stimulating electrode. An extracellular recording electrode (patch pipette filled with aCSF, as above) was placed at least 500 μm from the stimulating electrode, and paired pulse stimuli were used to confirm facilitation (average paired pulse ratio was 1.41±0.08 for ChR2-slices and 1.61±0.16 for ChR2+ slices). The stimulating current was adjusted until a minimum fEPSP (0.5 mV) was recorded extracellularly, and then reduced by half and kept constant for the rest of the experiment. A single current pulse was then delivered every 10 seconds to record a 5 minute baseline before the first plasticity induction protocol was delivered. The average baseline fEPSP was 0.28±0.08 mV in amplitude with 0.064±0.02 slope for ChR2-slices, and 0.24±0.05 in amplitude with 0.04±0.01 slope for ChR2+ slices.

Then the first plasticity protocol was delivered: the weak potentiating stimulus, which consisted of one electrical tetanus (i.e. one second of stimulation at 100 Hz). This typically leads to post-tetanic potentiation (PTP) where the fEPSP is briefly facilitated, decaying back to baseline between 30 sec to several minutes (Zucker & Regehr, 2002). We therefore tracked the fEPSP for 20 minutes after this protocol. The strong potentiating stimulus was then delivered, which consisted of four electrical tetani (one per second for four seconds). This typically leads to long-term potentiation (LTP), where the fEPSP remains facilitated well beyond 10 minutes (Bliss & Lomo, 1973). We therefore tracked the fEPSP for 30 minutes after this protocol. Finally, the strong depressing stimulus was delivered, which consisted of 900 single electrical pulses at 1 Hz. This typically leads to long-term depression (LTD) which plateaus around 10-15 minutes (Dudek & Bear, 1992), and we therefore tracked the fEPSP for a final 20 minutes after this protocol. The rising slope of the field potential was continuously monitored (0.1 Hz), and changes in its gradient were taken as an indication of synaptic potentiation (increased slope) or depression (decreased slope) (Bliss & Lomo, 1973 Figure 3).

Each of these plasticity induction protocols were interleaved with blue light stimulation (Figure 3 and supplementary Figure 1) in both ChR2-positive and -negative (control) slices. Specifically, 5 ms pulses of 470 nm light (10 mW) were delivered at 20 Hz for 30 seconds (designed to promote peptide release, Schöne et la., 2014), using a Lambda DG-4 fast beam switcher (Sutter Instruments) with a xenon lamp and ET470/40nm band pass filter, delivered through the 5× 0.1 NA microscope objective. Blue light was also delivered in this manner to examine the effects of MCH axon activation on spontaneous excitatory and inhibitory currents in hippocampal pyramidal cells (Figure 2).

### Fibre Photometry

Fibre optic implants were stereotaxically installed with the fibre tip above the lateral hypothalamus (angle 10°, bregma, −1.35 mm; midline, ±1.90 mm; depth, 5.10 mm) and fixed to the skull (25,28, 55). Fibre photometry was performed using the Doric fibre photometry system, in lock-in mode using simultaneous illumination with two LEDs (405-nm and 465-nm excitation, oscillating at 334 and 471 Hz, respectively; average power, ~100 μW at the fibre tip). Fluorescence produced by 405-nm excitation provided a real-time control for motion artefacts (Kim et al., 2016; Harris et al., 2022). To produce the plotted % ΔF/F values, the raw 405-nm–excited signal was fitted to the 465-nm–excited signal, then the % ΔF/F time series was calculated for each session as [100*(465 signal – fitted 405 signal)/fitted 405 signal], based on Lerner et al. (2015).

### T Maze

Each arm of the T Maze (Med Associates Inc.) measured 7 × 36.5 cm. The centre zone measured 9 × 9 cm. Mice (5 males and 8 females) spent the inter-trial interval (60 s) in a waiting chamber of 7 × 17 cm, from which they were allowed to enter the vertical arm of the maze at the beginning of each trial. The food cup zone was defined as a 3-cm wide perimeter around the food cup. The mouse was videotracked using Ethovision XT 15. Before the start of the experiment, mice were habituated to the maze with empty food cups, and to the liquid food reward (strawberry milkshake) in their homecage. Each learning session consisted of 12 trials, each of which had a maximum duration of 60 s. In correct trials, mice were returned to the waiting chamber after consuming the reward. In incorrect trials, a sliding door was used to confine the mouse in the incorrect arm of the maze for 10 s before returning it to the waiting chamber. Mice were trained on consecutive days until they reached the learning criterion of >=75% correct trials overall and >= ⅔ correct on the last 3 trials of the session. One food cup, which was kept in the same arm for each mouse, contained a 25 μL milkshake reward, while the other food cup was empty but had milkshake odour in order to prevent odour-based navigation. The maze was cleaned between each trial.

## Data Analysis

Statistical tests and descriptive statistics were performed as specified in Results and the figure legends. Data are presented as mean ± SEM, and a P value < 0.05 was considered to indicate significance. Analysis was performed using Synaptosoft, GraphPad Prism 8 and MATLAB R2019b (MathWorks).

## Author Contributions

J.J.H. and C.C. designed and performed the experiments and analysis, with input from D.P-G. and D.B. J.J.H. wrote the manuscript, with input from C.C., D.P-G. and D.B.

## Acknowledgements

We thank Olivia Shipton and Blanka R Szulc for advice regarding the hippocampal plasticity preparation, and Cornelia Schöne for advice regarding the optogenetic stimulation of MCH axons. This work was supported by the Francis Crick Institute (Burdakov Lab), which receives its core funding from Cancer Research UK (FC001055), the UK Medical Research Council (FC001055), and the Wellcome Trust (FC001055), by ETH Zürich, and by a Junior Research Fellowship from Imperial College London to J.J.H.

